# Screening Predictors of Weight Loss: An Integromics Approach

**DOI:** 10.1101/2020.07.06.188284

**Authors:** Joel Correa da Rosa, Jose O Aleman, Jason Mohabir, Yupu Liang, Jan L. Breslow, Peter R. Holt

## Abstract

Obesity has reached epidemic proportions in the United States but little is known about the mechanisms of weight gain and weight loss. Integration of “omics” data is becoming a popular tool to increase understanding in such complex phenotypes. Biomarkers come in abundance from high-throughput experiments, but small sample size is still is a serious limitation in clinical trials. It makes assessment of more realistic assumptions for complex relationships such as nonlinearity, interaction and normality more difficult. In the present study, we developed a strategy to screen predictors of weight loss from a multi-omics, high-dimensional and longitudinal dataset from a small cohort of subjects. Our proposal explores the combinatorial space of candidate biomarkers from different data sources with the use of first-order Spearman partial correlation coefficients. Statistics derived from the sample correlations are used to rank and select biomarkers, and to evaluate the relative importance of each data source. We tackle the small sample size problem by combining nonparametric statistics and dimensionality reduction techniques useful for omics data. We applied the proposed strategy to assess the relative importance of biomarkers from 6 different data sources: RNA-seq, RT-qPCR, metabolomics, fecal microbiome, fecal bile acid, and clinical data used to predict the rate of weight loss in 10 obese subjects provided an identical low-calorie diet in a hospital metabolic facility. The strategy has reduced an initial set of more than 40K biomarkers to a set of 61 informative ones across 3 time points: pre-study, post-study and changes from pre- to post-study. Our study sheds light on the relative importance of different omics to predict rates of weight loss. We showed that baseline fecal bile acids, and changes in RT-qPCR biomarkers from pre- to post-study are the most predictive data sources for the rate of weight loss.

## Introduction

Obesity has reached epidemic proportions in the United States, with about two-thirds of adults who are classified as being overweight or obese [10]. Little is known about the simultaneous effects of rapid weight loss induced by a clinically relevant very-low calorie diet (VLCD) on subcutaneous adipose tissue (SAT) inflammation, the plasma metabolome, fecal microbiome and bile acid content, and SAT transcriptome. It is well known that weight loss and weight gain occur at differing rates in individuals but the mechanisms responsible are still unclear [5, 16]. In obese subjects, gradual weight loss ameliorates adipose tissue inflammation and related systemic changes. A greater understanding of the factors that contribute to an individual’s enhanced rapid weight loss might greatly increase the development and efficacy of weight loss therapies.

Some existing gaps in disease understanding have been gradually filled by the development of omics and bioinformatics technologies and their ability to generate large amounts of data from biological systems. Omics science has been used for many goals; improving the accuracy of diabetes or heart disease predictive models, clustering cancers that were initiated by genetic aberrations, or revealing cell-type specific mechanisms in brain functions [3, 12].

Despite all the excitement around this so-called Big Data era, biomedical research is still is constrained by sample size restrictions. New research hypotheses are tested in a small number of subjects, preceding more extensive confirmatory studies. Although the costs for high-throughput experiments are decreasing, obtaining the omics profile for a large cohort is far from being feasible in clinical research.

The combination of high dimensionality, small sample size, and heterogeneous data sources, typical in multi-omics data, poses a challenge for analytical and inferential methods. This scenario prevents the use of classical statistical models and sophisticated machine learning algorithms. Assumptions such as normality, linearity, and interactive effects are difficult to be assessed; statistical tests are underpowered, and complex models cannot be externally validated.

Moreover, it is not feasible to implement the classic supervised learning paradigm of machine learning. Despite the fact that regularization techniques such as lasso-regression, ridge-regression and elastic nets [25] have been successfully used in situations where the number of features is much larger than the number of observations, assumptions such as linearity and additive effects are still difficult to be assessed with very small sample sizes. Kirpich and colleagues [15] pointed out that in multi-omics datasets, variable selection through these methods may inflate type-I error when handling small sample sizes.

The literature describing statistical methods to handle multi-omics data with small sample sizes is relatively new [18], but it still lacks parsimonious and informative solutions to unravel disease mechanisms.

## Materials and Methods

Our study proposes a method to screen predictors of an ordinal outcome in a multi-omics dataset obtained from a small cohort of subjects. An application of the proposed strategy utilized data from 6 different sources, examining the importance of each one in a study of the rate of weight loss in 10 obese individuals fed an identical diet in a metabolic ward [1]. Our pipeline is built to answer 3 scientific questions:

1. which are the most important weight loss predictors ?
2. which data sources show the highest predictive ability ?
3. how different data sources are interconnected when predicting the outcome ?

The small sample size issue is tackled by dimensionality reduction followed by the use of nonparametric statistics namely the first order partial Spearman correlation coefficient. P-values are not used as primary decision statistics but their regularity over a set of confounders is explored. We propose two decision statistics: the Biomarker Predictive Score (BPS) and the Persistent Significance (Ψ) that are jointly used to rank and select predictive biomarkers.

### Dataset

A prospective cohort study [1, 2] of VLCD-induced weight loss of 10% of baseline weight induced by an identical VLCD in ten grade 2-3 obese postmenopausal women studied in a metabolic unit collected data from 6 data sources : adipose tissue RNA-seq and RT-qPCR, plasma metabolomics, fecal microbiome, fecal bile acid and clinical data. Table 1 shows the number of biomarkers within each data source, the amount of missing data and the proportion of biomarkers in which the coefficient of variation is greater than 10%. Any biomarker with more than 20% missing data was excluded from the analysis.

**Table 1.** Biomarker data source frequency distribution.

| Data Sources | Total Variables<br>in Data Source | After Dimensionality<br>Reduction | > 20%<br>Missing Data | Biomarkers<br>with % CV > 10%<sup>1</sup> |
| --- | --- | --- | --- | --- |
| Clinical | 50 (0.11%) | 50 (12.14%) | 0 | 58% |
| Bile Acid | 24 (0.056%) | 24(5.83%) | 0 | 100% |
| RT-qPCR | 23 (0.054%) | 23 (5.58%) | 1 | 8.7% |
| RNA-seq | 41986 (98.78%) | 176 (42.72%) | 0 | 35% |
| Microbiota | 85 (0.20%) | 85(20.63%) | 0 | 0% |
| Metabolites | 336 (0.79%) | 54 (13.11%) | 0 | 57% |

The rate of weight loss (*WL*) is characterized as follows:

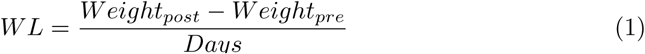

where *WL* is the daily average loss in kilograms observed in a study participant. To account for the fact that days on study were not uniform across subjects as well as the fact that the level of exercising is a potential confounder, we adjusted the daily rate of weight loss by the total number of steps during the study period. Therefore *WL*^*\**^ = *WL/*(Total Steps) was further used in the data analysis.

### Data Analysis

Our data analysis pipeline accomplished: a) dimensionality reduction; b) description of the amount of variability within each data source; c) assessment of the univariate association between weight loss rate and each one of the biomarkers; d) assessment of multivariate associations between weight loss rate and the biomarkers ; e) screening predictors of rapid weight loss; f) designing a network of interactions between data sources.

Our multivariate approach is described in Figures 1A-B where the use of low order partial rank correlation is the key element to rank predictors from different data sources. The strategy is also used to rank the data sources according to their relative importance when predicting weight loss rate.

**Figure 1.**
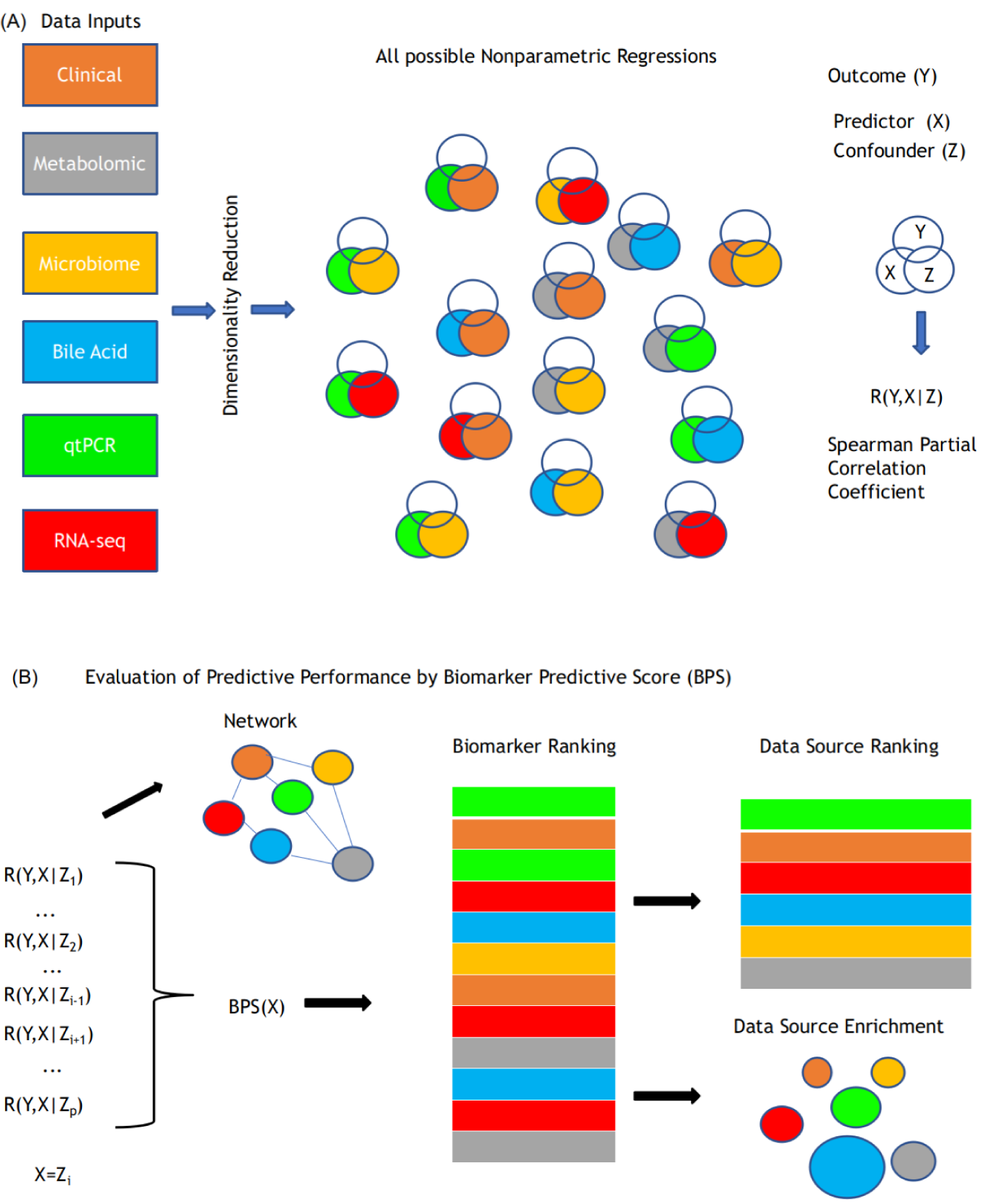
Schematic of multi-omic predictive model generation.. **(A)** Multiple data views serve as input to the predictive model. After dimensionality reduction, all possible regressions including two predictors are run. The rank first order partial correlation coefficient is used as a measure of association between Y and X after removing the effect of the potential confounder Z. **(B)** Evaluation of predictive performance. Partial correlation coefficients are computed over the confounders space and mapped to the BPS statistic. Biomarkers are ranked according to an importance metric, and this approach allows ranking and enrichment evaluation of data sources. A network based on aggregated measures of partial correlation is built to illustrate interactions between data sources.

### Correlation Analysis

Based on a sample of the bivariate random vector (*y, x*), our approach relies on the evaluation of the Spearman’s rank-order Correlation Coefficient in (2), a nonparametric measure of strength and direction of monotonic association between *y* and *x*:

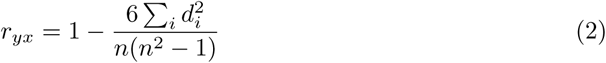

where *n* is the sample size and *d*_*i*_ is the difference between the ranks of *y*_*i*_ and *x*_*i*_. The Spearman correlation coefficient lies in the interval between −1 and +1, and the closer to the interval limits, the stronger is the evidence of association.

This coefficient is a less restrictive alternative to the Pearson product-moment correlation coefficient used to measure linear association between two variables. The only assumption used here is that all biomarkers are measured at least in an ordinal scale. Although this approach restricts the use of categorical variables, they can be included through the use of latent variables [8].

Our feature selection approach relies heavily on the first-order partial Spearman’s Correlation Coeficient described in equation (3),

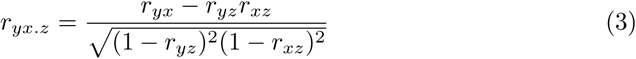

where *y* is the response variable, *x* is a biomarker and *z* is a potential biomarker confounder. This coefficient measures the monotonic association between *y* and *x* after adjusting, or accounting, for a potential confounding effect generated from a third variable *z*. An alternative way of evaluating this coefficient is by computing the correlation between the residuals from the regressions of *y* on *z* and *x* on *z* (see [14, 27]).

### Biomarker Predictive Score

Due to the small sample size restriction, we propose a metric for predictive importance that explores the partial correlation between the outcome and biomarkers from all data sources. We define our strategy as multivariate since all biomarkers are used in the construction of this metric. The use of the partial correlation coefficient for robust selection of features is described in [17] and also in [6] in an application to genomic data. The Biomarker Predictive Score evaluated according to (4),

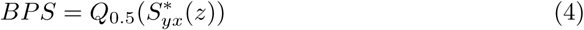

where *Q*_*p*_(*u*) is the *p*-th quantile of u’s distribution and 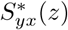, as function of *z*, is a scaled S-value [9], described in equation (5), obtained by combining the first-order partial Spearman’s correlation and its p-value.

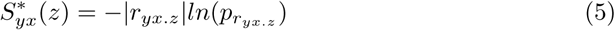

In (5), *r*_*yx.z*_ is the partial Spearman correlation in equation (3). 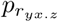 is the p-value derived from a hypothesis test based on the statistic in equation (6) described in [22, 26],

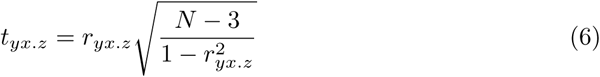

that follows the Student’s t distribution with N-3 degrees of freedom.

The partial correlation coefficient is measured across the set of confounders **Z**_−**j**_ that includes all biomarkers *z*_1_, *z*_2_, *…, z*_*p*_ except *z*_*j*_ = *x*, and tested for its significance for each one of the *p* − 1 elements of **Z**_−**j**_. This approach permits one to incorporate the magnitude of the association between *y* and *x*, its significance and confounding effects from all variables in the dataset.

The BPS statistic can be seen as a function of *p* − 1 random variables, assuming values in the positive real numbers. Consequently, its distribution is directly associated to the multivariate distribution of the random vector **Z**_−**j**_.

### Persistent Significance (Ψ_*α*_)

The BPS is useful for ranking the biomarkers according to their predictive ability but does not perform feature selection. To complement the BPS statistic in order to obtain a subset of important predictors, we propose the use of a complementary metric, the Persistent Significance Ψ_*α*_ in (7).

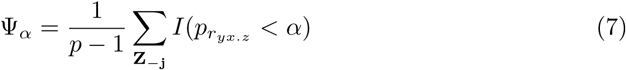

*I*(.) is an indicator variable. The statistic in equation (7) represents the proportion of times in which the confounding variable does not remove the partial correlation statistical significance, at a fixed *α* level. It is an index of robustness to the confounding in the association between *y* and *x*.

An ad hoc threshold *θ* for Ψ_*α*_ defines which biomarkers will be included in the set of selected features.

### Network Analysis

The partial correlation coefficient is further explored to assess how biomarkers interacts in a network when predicting the outcome. Considering *x* and *z* two vertices in a directed graph, with edges that are obtained according to the following metric :

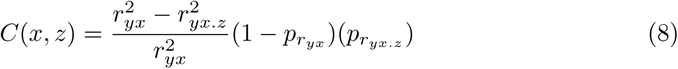

that can be seen as a measure of interaction between *x* and *z* when predicting the outcome *y*. In equation (8), *C*(*x, z*) represents the percentual decrease in the determination coefficient between *y* and *x* when accounting for a confounder *z*. As proposed in equations (5) and (7), the p-values from significance test are used as weights to leverage the connection strength between *x* and *z*. This metric accounts for the impact of the confounder *z* on the variability explained by the predictor *x*.

Consider different data sources labelled as *k ∈ {*1, 2, *…, K}*, in (9), we propose to evaluate the interaction between two data sources *k*_1_ and *k*_2_ with a graph with edges characterized as:

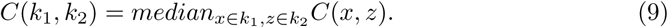

Based on *C*(*x, z*) and *C*(*k*_1_, *k*_2_), we applied weighted correlation network analysis as implemented in the R package qgraph [7] to build a network and to cluster predictors and data sources according to their similarity.

## Results

### Weight loss rates differ across individuals

Motivation for this application comes from data behavior displayed in Figures 2A-B. The rates of weight loss characterized by their longitudinal trajectories and estimated slopes differ markedly between study participants. In general, the participants showed a linear decay in weight at different speed rates. The addition of a quadratic term to describe the weight loss trajectory had no statistical significance.

**Figure 2.**
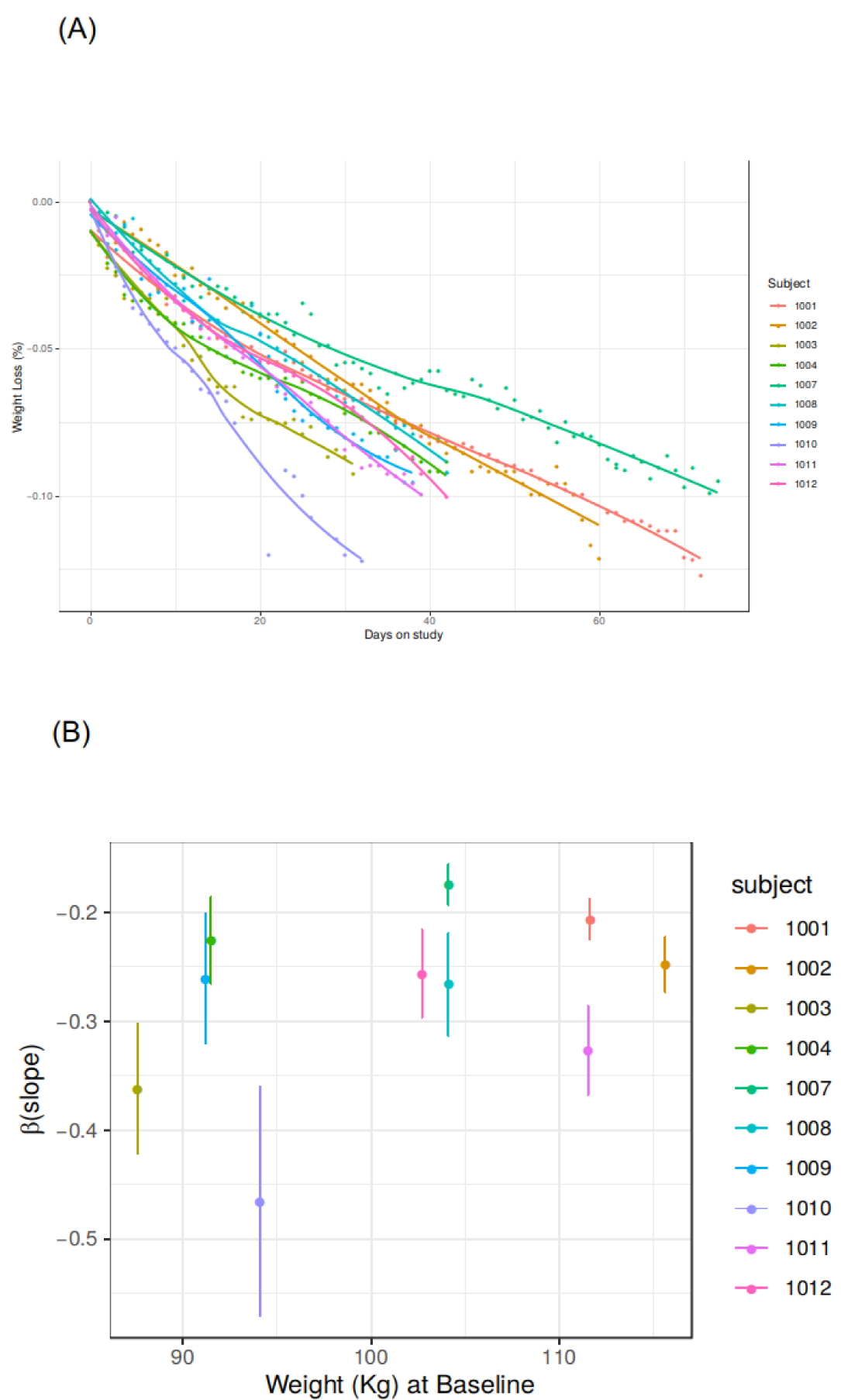
Rate of diet-induced weight loss is highly variable and not predicted by baseline weight.. **(A)**, Dynamics of the individual weight losses across the study period. The y-axis displays the % weight loss from baseline and x-axis the number of days after baseline at which the weight measurements were taken. Smoothed curves and their confidence intervals are estimated by the Loess method and overlaid to individual data points. **(B)**,Estimated slope coefficients and their 95% confidence intervals obtained from the OLS regression of the Weight Loss as a linear function of the days on study. The subjects are ordered according to their weights at baseline (x-axis).

The simplest predictor equation assuming each pound lost to be equivalent to 3300 kcal deficit did not predict individual rates. The slopes are different not only on their magnitude but also on the level of uncertainty estimated by the standard error. Although subjects were enrolled in different months over a period of one year, there is no statistical evidence of seasonal components affecting the weight loss dynamics.

To explain this variability, we explore information available in a wide format multi-omics data matrix containing a large number of features, but a small sample of subjects. The features are originated from different data sources: clinical, metabolomics, microbiome, fecal bile acids, RNA-seq and RT-qPCR gene expressions. The strategy for screening potential predictors of weight loss is carried out by assembling information gathered from simple statistical models rather than using complex formulations.

### Dimensionality Reduction

The large number of individual genes analyzed in the adipose tissue RNA-seq and plasma metabolite datasets required dimensionality reduction from these sources. To prevent a selection bias due to this problem, the dimensions in RNA-seq and metabolites biomarkers were reduced by mapping gene expressions and metabolites concentration to pathways scores, evaluated for each subject. The Gene Set Variation Analysis (GSVA, [11]) as implemented in the gsva R package was applied to reduce the large number of RNA-seq genes and metabolites to a manageable number of pathways. GSVA scores were computed for 4107 canonical pathways curated from online databases and available at Broad Institute website. An additional step was carried out to reduce this number to 176 pathways gene sets, selected according to the nominal significance of their estimated differences from pre- to post-VLCD with a mixed-effects model. The set of 336 metabolites was mapped to a reduced set of 87 metabolic pathways.

### Data source variability

The data sources are heterogeneous, both in their between- and within-individuals variability. For predicting weight loss, large between-individuals variability in biomarkers can be beneficial to discriminate the most predictive ones. On the other hand, large within-individual variability may not add much gain to a rank-based method since a subject may be ranked differently at different time points. To establish a fair comparison between data sources, we have evaluated for each biomarker the coefficient of variation at baseline, post-VLCD, and the changes from pre- to post-VLCD. Figure 3A boxplots summarize the variability with the log-scaled coefficient variation in biomarkers from different data sources.

**Figure 3.**
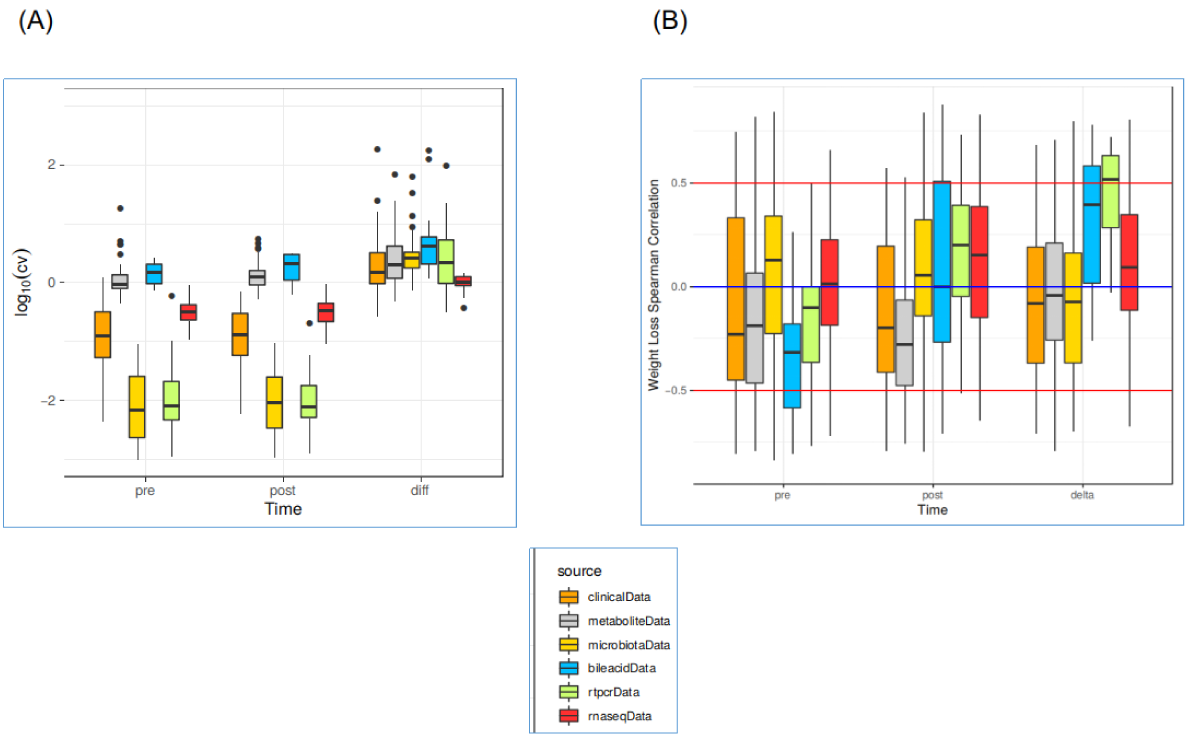
Bile acids and gene expression by RT-qPCR are most informative before and during weight loss, respectively. **(A)** The boxplots illustrate variation in measurements before (pre), after (post) or during (diff) weight loss according to the different data sources and time references. The boxes are built based on the quartiles of absolute coefficient of variation in log-scale. Negative values are associated with lower variability in the biomarkers as shown by microbiota and RT-qPCR pre and post study. **(B)** The boxplots show the correlation of biomarkers with weight loss depends on the data source and time point in the study. The box shows quartiles of the Spearman Correlation Coefficient. The horizontal blue line locates the null correlation, and red lines correlations of intermediate magnitude (*r* = 0.5). Proximity of the center of the box to the red lines shows that bile acids at pre study as the most highly correlated data source, and metabolites at post measurements. Changes (diff) demonstrate RT-PCR as the most highly correlated data source.

At each time point, pre- and post-VLCD, bile acids and metabolic pathways exhibit the most extreme biological variation, while gut microbiome and RT-qPCR have the least extreme ones. The low biological variation in microbiota biomarkers is partially due to the excess of zeros, a common problem when handling this kind of data [13]. The same applies to RT-qPCR where several measurements are close or below the lower limit of detection (LLOD).

The coefficient of variation expectedly rises in all data sources when evaluating changes from pre- to post-VLCD, at a less extent in RNA-seq, fecal bile acids, and metabolites. The variance computed for the difference between two variables is reduced when these two variables are positively correlated (see [20]). Therefore, the small increment suggests high, and positive, correlation between pre- and post-VLCD measurements within these three data sources.

### Biomarkers Correlation with Weight Loss grouped by Data Source and Time Point

Figure 3B illustrates how the correlation between weight loss rate and biomarkers ranges across data sources and different time points. In this figure, the blue horizontal line is placed at the null association level(*r*_*yx*_ = 0), and the red horizontal lines represent moderate association (*r*_*yx*_ = 0.5) according to the Spearman correlation coefficient. RT-qPCR changes from baselines are the unique scenario in which the median Spearman correlation coefficient exceeds moderate correlation levels with weight loss rate. Bile acids’ correlations are close to this level but still below the 0.5 threshold.

### Biomarker and Data sources Ranked and Selected According to Predictive Importance and Persistent Significance

The multivariate approach ranks and selects the biomarkers according to the proposed Biomarker Predictive Score (BPS) and persistent significance metrics. After ranking, our feature selection criteria extracts a robust subset of predictive biomarkers by using the persistent significance Ψ_*α*_ as defined in equation (7). Two ad hoc hyperparameters need to be defined when using this statistic, the significance level, fixed at the classic 5% level and the persistence threshold that represents the proportion of correlations that remain significant after adjusting for all possible confounder from all different data source. This threshold was set to 70% and whenever Ψ_0.05_ *<* 0.70 the biomarker was included in the final set.

The panel in Figures 4A,B summarizes the results of the application of this pipeline. Figure 4A shows a bar plot with the BPS statistic estimated for biomarkers selected in each time point, considering a persistent significance greater than 0.7. In pre-, post-VLCD and changes from baseline, the procedure selected 26, 27 and 22 biomarkers, respectively (see Supplementary Table S1).

**Figure 4.**
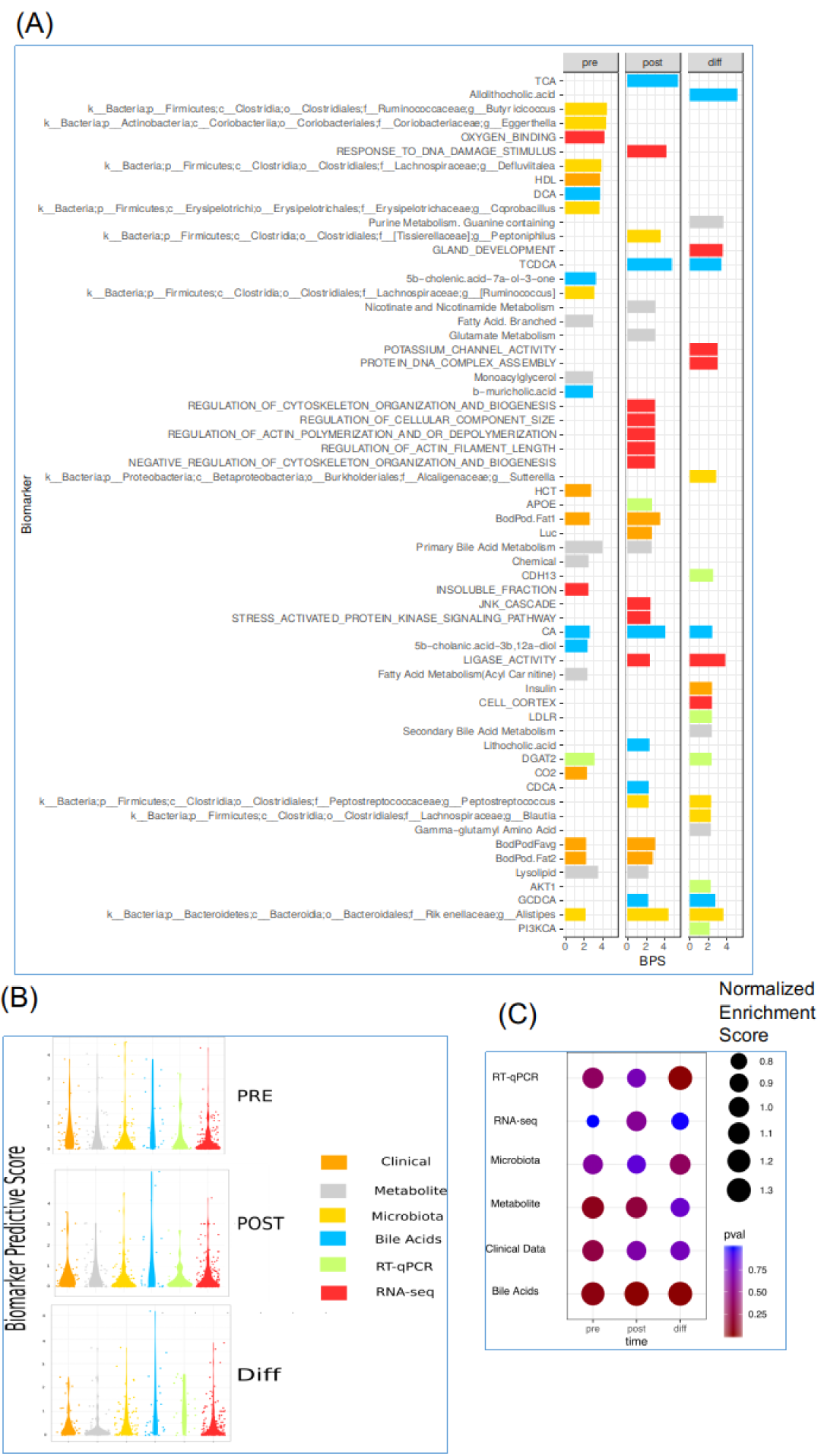
Biomarker prediction of weight loss rate. **(A)** Biomarkers selected at pre-study, post study and during (diff) study according to BPS and persistent significance *>* 0.7. Biomarker description is labelled in the Y-axis and colors discriminate the data source. **(B)** Violin plots show BPS distribution for different data sources at pre-study, post study and during (diff) study. **(C)** A bubble plot based on Gene Set Enrichment Analysis (GSEA) results explains how data sources are enriched in the ranked list of biomarkers. The ranking is built according to the BPS statistic. The size of the bubble is proportional to the normalized enrichment score and the gradient color schema indicates statistical significance. Bile acids are significantly enriched in all time points and RT-qPCR data source has largest enrichment in during (diff) study.

Among the biomarkers selected from changes from pre- to post-VLCD, the most represented data sources are RNA-seq (22.8%) and RT-qPCR (22.8%) followed by fecal bile acids(18.2%) and fecal microbiota (18.2%), then metabolites(13.6%) and only one (4.5%) clinical biomarker. In terms of relative overrepresentation, fecal bile acids and RT-qPCR are the most significant ones. When taking into consideration the ranking, fecal bile acids overperform RT-qPCR markers.

In Figure 4B, violin plots show the distribution of BPS statistic across data sources and time points. The different distributions suggest that data sources have different degrees of importance in the weight loss prediction. The central tendency of the distribution is a proxy for the data source importance and the dispersion indicates how heterogeneous is the information contained in this specific block of data. These figures also show that the importance of data sources is relative to the time reference pre-VLCD, post-VLCD or Δ). The densities in Figure 4B are skewed towards zero. A high concentration of biomarkers close to zero indicates that most of them have no impact in explaining variation in weight loss rates.

### Enrichment of Data Sources

The multivariate correlation analysis and the resulting BPS provide a way of ranking all biomarkers independent of which data source the biomarker comes from. To verify the enrichment of the data sources on top ranked biomarkers, we applied the Gene Set Enrichment Analysis (GSEA) [24] on a pre-ranked list of biomarkers sorted in descending order from the highest to the lowest BPS. The results are presented in Figure 4C. This analysis can be seen as an indirect way of quantifying the predictive importance and determining the statistical relevance of each data source. The Normalized Enrichment Score was used as statistic for enrichment and p-value for its significance.

### RT-qPCR and Fecal Bile Acids as Network Hubs

Lastly, we aimed to understand the connectivity between the examined biomarkers by network analysis. In Figure 5A-C we show 3 networks built with the connectivity measures described in equations (8) and (9) for each time point; pre-, post-VLCD and for the Δ changes from pre- to post-VLCD. The directed arrows in the network indicate how a data source in the origin node impact the weight-loss predictive capability of the data source in the descendant node. In Figure 5C, all data sources have a clear impact on RT-qPCR, meaning that the relationship between the gene expression and the rate of weight loss is largely modified when the partial correlation adjusts for a biomarker from a specific data source (e.g. microbiota). An expected self-loop in RT-qPCR is also expected since the selected genes for amplification are supposed to be jointly linked to the weight loss rate. This dynamic is different from the one in Figure 5A where fecal bile acids, and clinical data sources are most affected by RT-qPCR and RNA-seq. Clinical data at baseline has a larger impact from RNA-seq and microbiota. Figure 5B shows that the network built based on post-VLCD data has as configuration that is similar to baseline (pre-VLCD), except for the fact that fecal bile acids hub position is not as evident.

**Figure 5.**
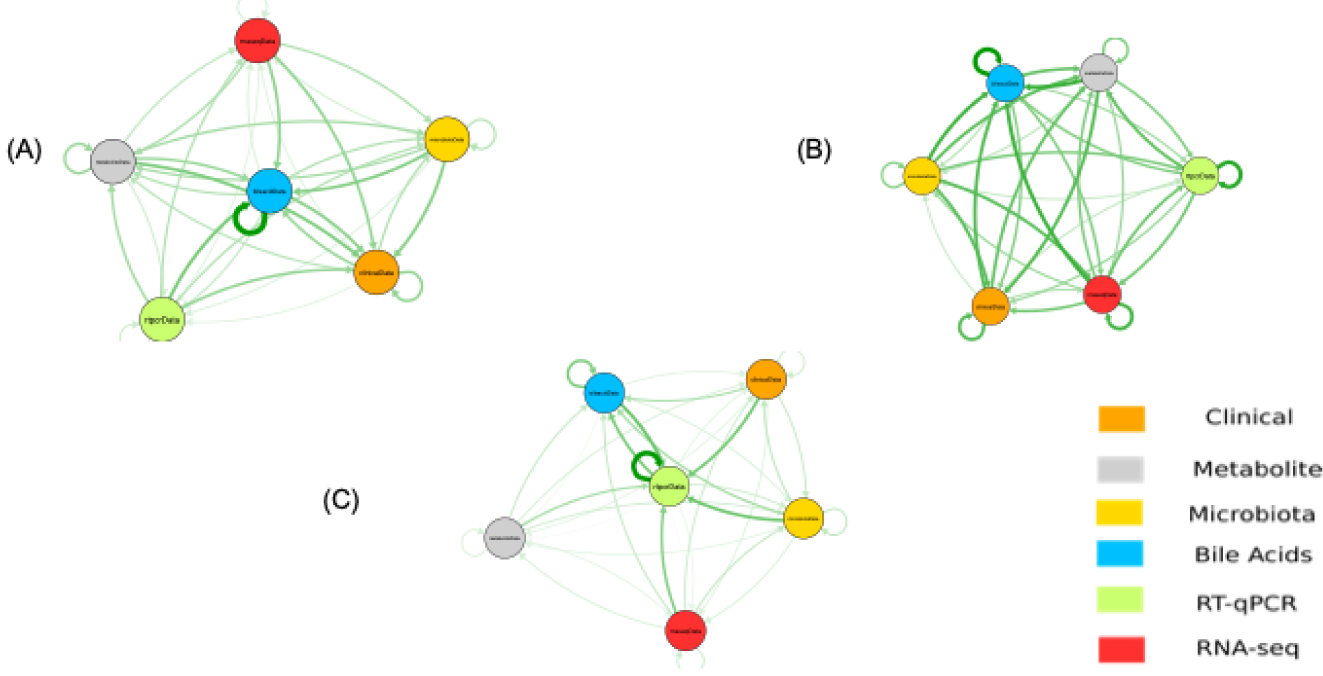
Network analysis of biomarker interactions for weight loss rate. Networks analyses at pre-study, post-study, and changes from pre- to post-study (diff) show how data sources interact when predicting the weight loss rate. Nodes represent individual data sources, and the line thickness represents the strength of the interaction. **(A)** Pre-study bile acids interact with genomic data sources (RT-qPCR and RNA-seq). **(B)** Post-study network preserves the structure in (a) where bile acids interact with higher intensity with RT-qPCR, RNA-seq, and, to a lesser extent, with microbiota and metabolites. **(C)** Diff network structure shows RT-qPCR occupying a hub position, interacting with other data sources.

## Discussion

We propose a method for ranking and selecting predictive biomarkers in a high-dimensional study with a small sample size. A multi-omics pipeline integrates different data sources, identifying how they are interconnected. The strategy relies heavily on the use of the first-order partial correlation between the outcome and potential predictive biomarkers. Two metrics BPS and the PS score, are combined to generate a final set of predictive biomarkers.

This methodology is applied to identify predictors of weight loss rate in a small cohort of subjects provided a very low-calorie diet. We used this strategy in the pre-study, post-study, and changes from pre to post study (diff) timepoints for each data source.. Before integrating data derived from multiple sources, we investigated factors that contributed to their heterogeneity. Fecal bile acids, for example, show pre-study greater biological variability but also greater magnitude in their correlation with the rate of weight loss when compared to post study data (see Figure 3A). Unlike other data sources that are heavily skewed towards lower BPS values, fecal bile acids are over-represented in the upper tail of the BPS distribution (see Figure 3B).

Finally, in Figure 5A, for prestudy data bile acids occupy a hub position in the interaction network. In the diff data, fecal bile acids and RT-qPCR biomarkers also are an essential predictive data source, especially when investigating changes from pre to post-study.

Different approaches confirm the importance of fecal bile acids and RT-qPCR as predictors of weight loss, such as the, shifted distribution in the univariate correlation displayed in Figure 3A, the BPS distribution in Figure 4, and also the interaction network, as shown in Figure 5C. Among the 22 markers selected for diff, five came from the RT-qPCR data source. Gene Set Enrichment Analysis (GSEA) shows significant enrichment of bile acids and RT-qPCR in the top positions on the list of biomarkers ranked according to predictive power. The fecal bile acid is enriched at all 3 time points and RT-qPCR from pre to post-study(Figure 4C). Even though RNA-seq is the data source with the largest number of predictors (176 pathways), it shows weak enrichment in the top predictive group. Although metabolic pathways show similar biological variation as fecal bile acids (see Figure 3A), their association with weight loss is only evident in the pre-study data, as seen in Figures 3B, 4A,C. Their minor contribution is further demonstrated by their position in the interaction network at the periphery with weak connections (Figures 5A,C). Overall, our data shows that clinical, microbiota, and RNA-seq data sources are of less predictive importance. Microbiota only appear in the list of 5 top predictive markers pre study but not at other times. Clinical data were not found to be as important as other sources and their changes from pre to post-study are under-represented in the list of selected predictive biomarkers (see Figure 4A).

## Conclusion

The present study was launched by the observation that a group of 10 obese postmenopausal women placed on an identical very low calorie diet in a metabolic facility lost weight at rates that varied by over 100%.Major variations in weight loss [23] and weight gain [4] rates have been observed in previous studies. Hypotheses about the determinants of weight loss rates in individual subjects have focused on genetic factors, differences in metabolic rates, energy expenditure. microbiota composition and metabolites. Some individuals have been shown to have a “thrifty” phenotype, with lower weight loss rates during calorie restriction than those with a “spendthrift” phenotype, who lost more weight [21].

Our study attempted to examine the importance of several potential biomarker sources on a predictive model of the rates of weight loss. Striking was data that showed the impact of fecal bile acids on predicting weight loss rate from the pre-study data as well as the difference between pre and post study results. Bile acids in the gut are now recognized as powerful signaling molecules for receptors that act on systemic lipid and carbohydrate metabolism [19]. Further studies on the biologic role of gut bile acids in determining the rate of weight loss in individuals during caloric restriction are warranted.

## Abbreviations

BPS: Biomarker Predictive Score
GSEA: Gene Set Enrichment Analysis
GSVA: Gene Set Variation Analysis
RT-qPCR: Real Time quantitative Polymerase Chain Reaction
SAT: Subcutaneous Adipose Tissue
VLCD: Very Low Calorie Diet
WL: Weight Loss

## Declarations

### Ethics approval and consent to participate

The study was approved by the Institutional Review Boards at Rockefeller University, Weill Cornell Medical College, and Memorial Sloan Kettering Cancer Center, and registered under ClinicalTrials.gov identifier NCT01699906.

### Consent for publication

Not applicable.

### Availability of Data and Materials

Clinical Trial design is reported at NCT01699906. Deidentified transcriptomic data is available at GEO Accession Number GSE106289

### Author’s contributions

JCR, JOA, JLB and PRH conceived the study, JCR and JM performed the statistical design and analyses. JOA executed the clinical, transcriptomic and RT-PCR analyses, YL executed bioinformatics analyses, JCR, JOA, JLB and PRH wrote the manuscript with input from all coauthors.

## Acknowledgements

We thank the participants of the original clinical study and the coauthors of prior publications from this study.

